# Expanding the language network: Domain-specific hippocampal recruitment during high-level linguistic processing

**DOI:** 10.1101/091900

**Authors:** Idan A. Blank, Melissa C. Duff, Sarah Brown-Schmidt, Evelina Fedorenko

## Abstract

Language processing requires us to encode linear relations between acoustic forms and map them onto hierarchical relations between meaning units. Such relational binding of linguistic elements might recruit the hippocampus given its engagement by similar operations in other cognitive domains. Historically, hippocampal engagement in online language use has received little attention because patients with hippocampal damage are not aphasic. However, recent studies have found that these patients exhibit language impairments when the demands on flexible relational binding are high, suggesting that the hippocampus does, in fact, contribute to linguistic processing. A fundamental question is thus whether language processing engages domain-general hippocampal mechanisms that are also recruited across other cognitive processes or whether, instead, it relies on certain language-selective areas within the hippocampus. To address this question, we conducted the first systematic analysis of hippocampal engagement during comprehension in healthy adults (*n*=150 across three experiments) using fMRI. Specifically, we functionally localized putative “language-regions” within the hippocampus using a language comprehension task, and found that these regions (i) were selectively engaged by language but not by six non-linguistic tasks; and (ii) were coupled in their activity with the cortical language network during both “rest” and especially story comprehension, but not with the domain-general “multiple-demand (MD)” network. This functional profile did not generalize to other hippocampal regions that were localized using a non-linguistic, working memory task. These findings suggest that some hippocampal mechanisms that maintain and integrate information during language comprehension are not domain-general but rather belong to the language-specific brain network.

**Significance statement:** According to popular views, language processing is exclusively supported by neocortical mechanisms. However, recent patient studies suggest that language processing may also require the hippocampus, especially when relations among linguistic elements have to be flexibly integrated and maintained. Here, we address a core question about the place of the hippocampus in the cognitive architecture of language: are certain hippocampal operations language-specific rather than domain-general? By extensively characterizing hippocampal recruitment during language comprehension in healthy adults using fMRI, we show that certain hippocampal subregions exhibit signatures of language specificity in both their response profiles and their patterns of activity synchronization with known functional regions in the neocortex. We thus suggest that the hippocampus is a satellite constituent of the language network.

## Introduction

The hippocampus has long been recognized as a key component of the declarative memory system (1, 2). It plays a critical role in binding together information from multiple sources across time and space in order to generate a representation that relates the co-occurring aspects of an experience to one another: a relational memory (3, 4). This ability to keep track of the relations among entities and events is fundamental across cognitive domains, from understanding causality, to remembering locations of objects in space, to learning new words. Therefore, diverse cognitive functions may require hippocampal contributions. Indeed, hippocampal pathology has been linked to poorer behavioral outcomes across a range of cognitive and social domains (5). Here, we focus on the domain of language and ask: how does the hippocampus contribute to language processing?

Online language use appears to place critical demands on relational representation and processing (6): it requires mapping the linear relations between acoustic forms to hierarchical relations between units of meaning, across multiple levels of granularity (from morphemes to sentences, 7); keeping track of semantic relationships across multiple time-scales (from rapid, incremental processing to extended discourse use, 8); and integrating information across modalities (e.g., incorporating contextual information, eye-gaze or speech-accompanying gestures, 9, 10-15). Traditionally, however, hippocampal contributions to online language use have not received serious consideration, as decades of research on patients with acquired hippocampal damage have established that such patients are not aphasic despite their profound impairments in relational memory (16–18). Instead, research has focused on the role of the hippocampus in the acquisition of vocabulary, i.e., establishing and maintaining memories of the arbitrary relations between word forms and their meanings (19, 20). This focus on word learning is in line with the widespread view that the hippocampus contributes exclusively to relational representations that endure over time (long-term memory) and not to information that is temporarily stored and manipulated during ongoing behavior (working memory) (1, 3, 21). Because psycholinguistic theories of online processing typically focus on the latter form of memory (22–29), the hippocampus has not been considered part of the language network.

Nonetheless, two sources of evidence suggest that we should reconsider potential hippocampal contributions to language processing. First, accumulating neuropsychological (30, 31) and neuroimaging (32, 33) evidence is challenging the view that hippocampal involvement in relational binding is limited to long-term memory. Instead, relational representations appear to be hippocampus-dependent even when maintenance is only required for a few seconds, on the time-scale of working memory. Thus, hippocampal representations are available early enough to guide ongoing behavior and can be flexibly expressed when relational information needs to be manipulated online. Thus, the hippocampus may also process relational information during online language use (6).

Second, even though damage to the hippocampus does not result in aphasia, it still confers deficits in real-time language processing when the demands on flexible relational binding are high. For instance, individuals with bilateral hippocampal damage exhibit compromised comprehension when they need to maintain and integrate recently available information in order to interpret ambiguous expressions that refer to one of two competing discourse referents (34, 35). Similarly, in language production, these individuals are less likely to mark previously established referents as “shared knowledge” during conversation (using, e.g., “a man” instead of “the man”; 36), even though they can acquire novel, self-generated labels for such referents and maintain these over months (37). More generally, individuals with amnesia produce language with limited flexibility, including relatively infrequent combinations of different communicative devices, e.g., task-related talk with verbal play (38), direct communication with reported speech (39), or speech with gesture (40). Thus, at least some aspects of language processing appear to depend on the relational representations supported by the hippocampus (cf. 41, 42).

Whereas current evidence strongly suggests that the hippocampus contributes to online language use, the nature of its contributions remains unclear. Perhaps the most fundamental question is whether the hippocampus supports only domain-general relational processes that are crucial for language but are not language-selective, or whether instead some parts of the hippocampus are more tightly integrated specifically with the language processing system. On the one hand, language is a relatively recent artifact of human-unique cultural evolution (43) and the hippocampus is an evolutionarily older structure conserved across species (44). Thus, hippocampal mechanisms that support language processing may not seem a priori likely to be language-specific. On the other hand, functional specificity can - and does - emerge as a function of our experience with the world (45–47). Given the importance of language in our lives and the evidence for a neocortical language network that is highly domain-specific, at least in adults (48), it is possible that parts of the hippocampus become selectively coupled with the language network and show similar functional signatures.

To address this question, as well as complement existing patient studies, we use fMRI to functionally characterize hippocampal engagement in online language processing in healthy individuals. Specifically, we first tested whether we could functionally localize language-responsive sub-regions within the hippocampus that would be recruited reliably and robustly during language use (Experiment 1). Then, we tested whether such regions respond to language processing selectively or, alternatively, also respond to non-linguistic tasks that place high demands on working memory, flexible mappings, and relational binding (Experiment 2). Finally, we tested whether such regions systematically and selectively co-activate with the neocortical language network (49–54), especially with increasing demands on language processing (Experiment 3).

## Results

A total of *n*=150 individuals participated in the study with different, partially overlapping subsets participating in the three experiments, as detailed in the following sections. For all experiments, candidate language-responsive sub-regions in the two hippocampi were identified individually in each participant using a comprehension task that contrasted sentences and lists of unconnected, pronounceable nonwords (Fig. 1A). Unlike nonword reading, which involves perceptual and phonological processing but no higher-level linguistic processes, sentence reading entails accessing word entries in the mental lexicon and, critically, relational processing: semantic and syntactic composition of words and phrases. Therefore, hippocampal sub-regions recruited during linguistic processing should respond more strongly to sentences than to nonwords. The sentences > nonwords contrast has been previously demonstrated to activate neocortical regions that comprise the language network (49, 55).

**Figure 1.**
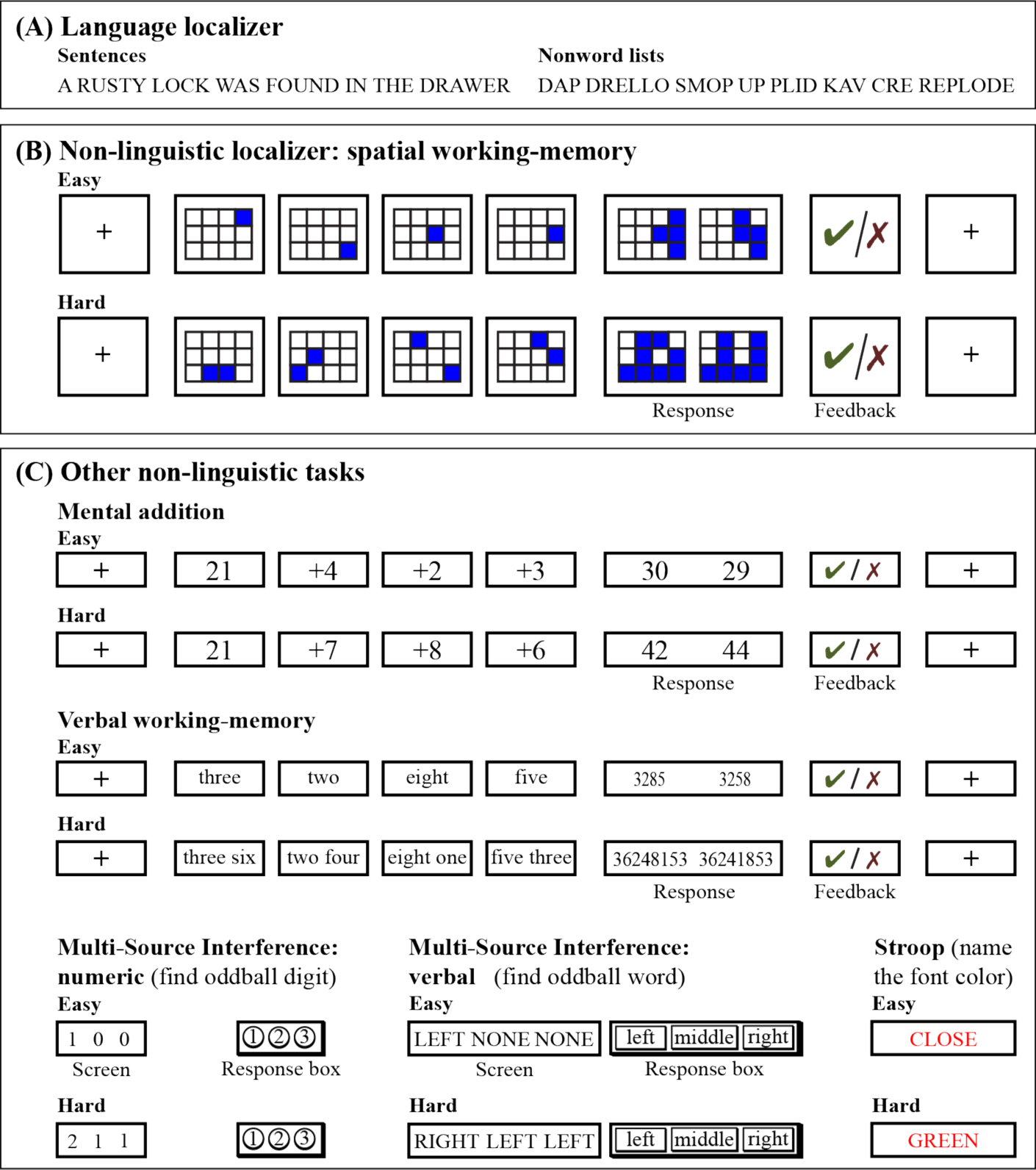
Materials and tasks. (A) Sample stimuli from the comprehension task used to localize language-responsive fROIs. (B) Sample trials in the easy and hard conditions of the non-linguistic task used to localize control fROIs. (C) Sample trials in the easy and hard conditions for each of the five non-linguistic tasks used in Experiment 2.

To test whether the functional characteristics of these functional regions of interest (fROIs) were unique within the hippocampal neural milieu, we also identified sub-regions that responded, above a fixation baseline, to a non-linguistic, working memory task requiring relational processing of spatial locations (Fig. 1B). This task does not recruit the neocortical language network (55), so if the language-responsive hippocampal sub-regions are domain-specific they should similarly not be recruited by this task. Instead, this task would localize a different, functionally distinct sub-region (We note that in Experiments 2 below, for which data on this task were not collected, we instead identified sub-regions that responded above fixation baseline to reading nonwords in the language localizer task. In the neocortex, this contrast recruits regions outside the language network similar to those identified by the spatial working-memory task; e.g., 56).

Both kinds of hippocampal fROIs (language-responsive and control) showed a consistent topography across participants (*n*=93, the sample from Experiment 1) (Fig. 2). Their centers were separated from each other by 13.4mm on average (SD 7.1) in the left hemisphere (LH) and by 12.8mm (SD 6.9) in the right hemisphere (RH). To characterize their topography, we computed for each participant the Center of Mass (CoM) of each fROI as well as the CoM of the entire hippocampus. We then projected these locations onto two orthogonal axes describing the shape of the hippocampus: an axis extending from its posterior-superior end to its anterior-inferior end, and a medial-lateral axis. Language-responsive fROIs were consistently located anterior-inferior to the center of the hippocampus, in both the LH (*p*<10^−22^, Cohen’s *d*=1.46) and the RH (*p*<10^−12^, *d*=0.94). In contrast, control fROIs were posterior-superior to the center of the hippocampus (LH: *p*<10^−8^, *d*=0.73; RH: *p*<10'^u^, *d*=0.88) and thus differed from language-responsive fROIs (LH: *p*<10^−22^, *d*=1.43; RH: *p*<10^−18^, *d*=1.23). Furthermore, in the LH both language-responsive and control fROIs were located medial to the center of the hippocampus (*p*<10^−4^, *d*=0.48 and *p*=0.001, *d*=0.39, respectively), but they did not differ from one another on this axis (*p*=1, *d*=0.07). In the RH, only the language-responsive fROI showed this medial bias (*p*<10^−8^, *d*=0.75); it differed from the control fROI (*p*<0.001, *d*=0.45), which did not show this bias (*p*=0.44, *d*=0.16) (here, and in each of the following experiments, significance tests are FDR-corrected for multipel comparisons, 57).

**Figure 2.**
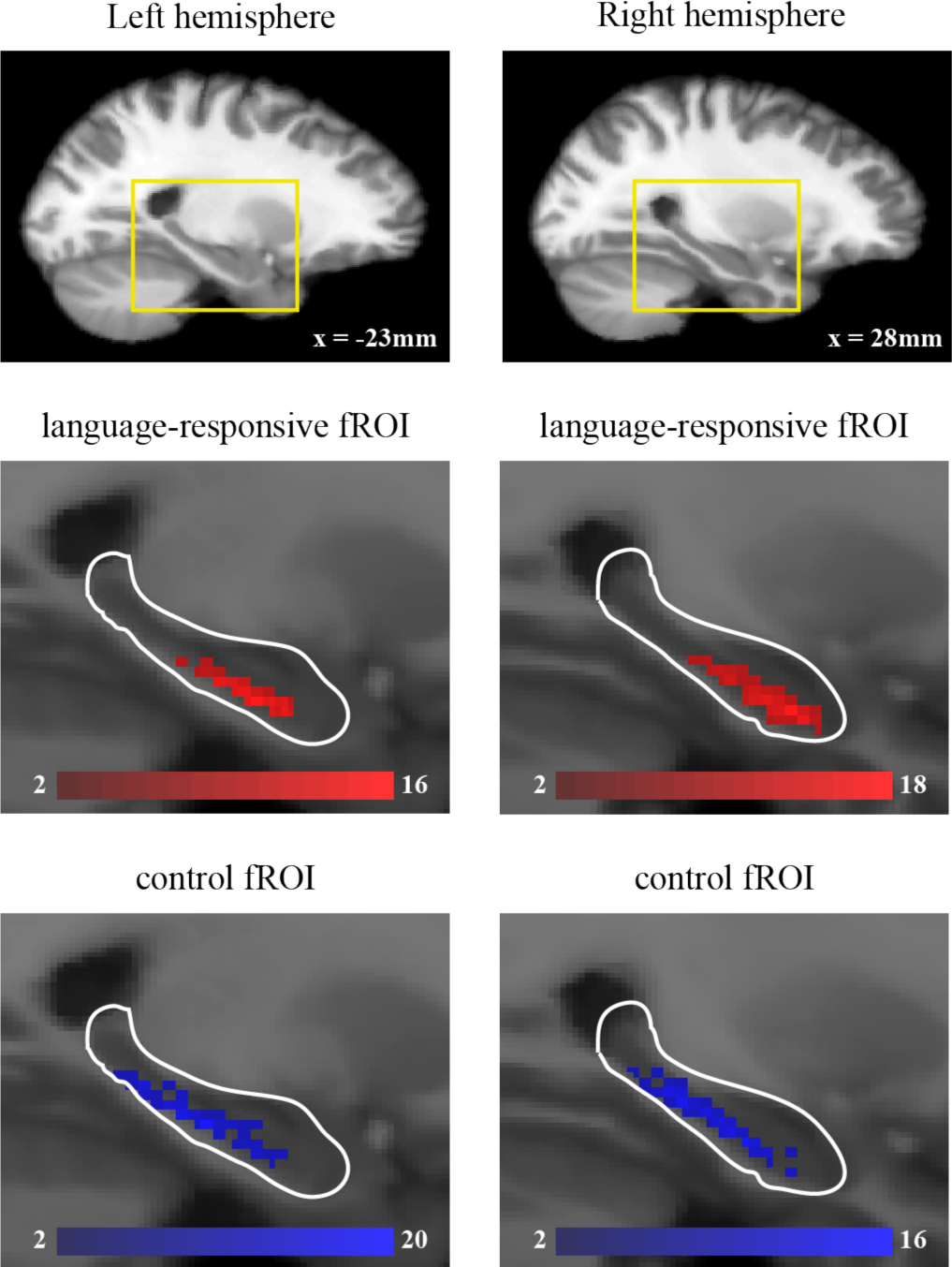
Topography of hippocampal fROIs. Top: a sagittal brain slice is shown for each hemisphere (column), taken from Freesurfer’s average T1 image in the common (MNI) space. A segment of the slice in which the hippocampus is visible is bounded in yellow, and is shown enlarged in the images below. Middle: the distribution of language-responsive fROIs across participants. Each red-colored voxel shows the number of participants whose fROI has its center of mass in that location (collapsed across the medial-lateral axis). The contours of an anatomical mask of the hippocampus are shown in white; note that the locations of language-responsive fROIs within this mask are relatively anterior. Bottom: the distribution of control fROIs. Conventions are the same as for the middle panel.

### Experiment 1: Language-responsive fROIs are reliably and robustly recruited during language processing

We estimated the responses of language-responsive (*n*=90 participants) and control fROIs (*n*=71) to each condition of both localizer tasks using across-run cross-validation (58) (Fig. 3A). During the comprehension task, language-responsive fROIs responded significantly to sentences (LH: *p*=0, *d*=0.8; RH: *p*<0.001, *d*=0.4) but not to nonwords (LH: *p*=0.17, *d*=0.21; RH: **p*=1*, *d*=0.05), with a significant difference between the two conditions (LH: *p*<10^−13^; *d*=0.9; RH: *p*<10^−6^, *d*=0.61). Therefore, these fROIs exhibit a fundamental functional signature characteristic of language processing: a greater response to meaningful and structured linguistic representations than to meaningless and unstructured ones.

**Figure 3.**
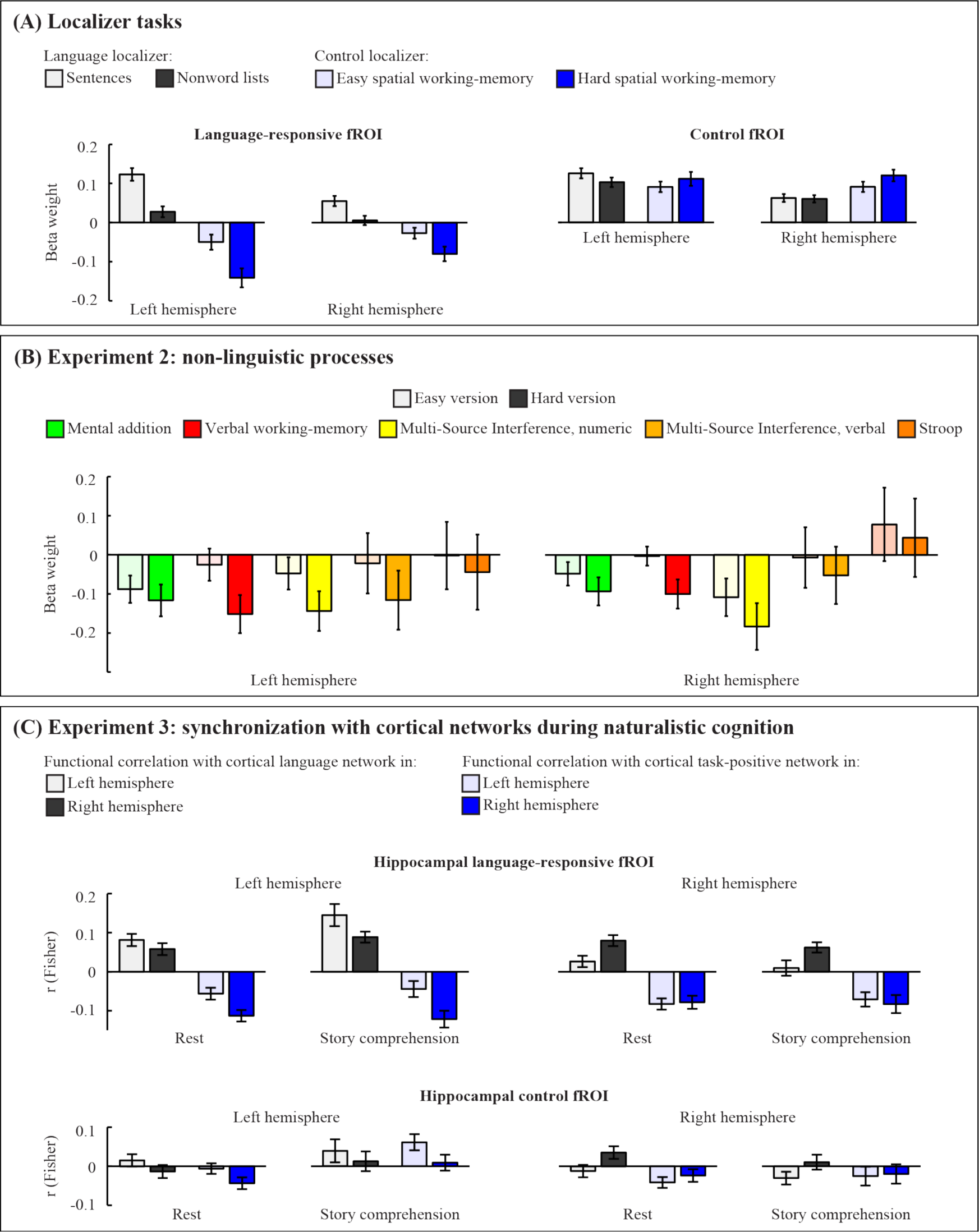
Functional profiles of language-responsive and control fROIs in the hippocampus (for details, see Results). For figures (A)-(B), data used to define fROIs and data used to estimate response magnitudes are independent.

In contrast, control fROIs responded to reading both sentences (LH: *p*=0, *d*=1.03; RH: *p*=0, *d*=0.65) and nonwords (LH: *p*=0, *d*=0.91; RH: *p*=0, *d*=0.67). Surprisingly, the LH control fROI showed a small linguistic Sentences > Nonwords effect (*p*=0.003, *d*=0.37; RH: *p*=1, *d*=0.035). Critically, however, language-responsive fROIs exceeded their ipsilateral control fROIs in the size of this linguistic Sentences > Nonword effect (LH: *p*<10^−12^; *d*=0.87; RH: *p*<10^−10^, *d*=0.76). Moreover, in language-responsive fROIs, the Sentences > Nonwords effect showed a bias towards the left (language dominant) hemisphere (*p*<10^−4^, *d*=0.49), an effect that distinguished these fROIs from control fROIs (*p*=0.002, *d* = 0.38) which did not show such bias (*p*=1, *d*=0.07).

Unlike language processing, performing a spatial working memory task deactivated the language-responsive fROIs when the task was hard (LH: *p*=0, *d*=0.62; RH: *p*=0, *d*=0.45), indicating that relational processing does not recruit these regions equally across domains; instead, these fROIs might be language-specific (see Experiment 2). The easy version of the task did not cause any change in activity relative to fixation (LH: *p*=0.09, *d*=0.27; RH: *p*=0.24, *d*=0.2), leading to an Easy > Hard effect (LH: *p*<10^−10^, *d*=0.78; RH: *p*<10^−5^, *d*=0.54), similar to what has been reported for some of the neocortical language regions (55). The Easy > Hard effect too showed a LH bias (*p*=0.005, *d*=0.34) not different in size from the bias of the linguistic effect (*p*=1, *d*=0.03).

In control fROIs, the non-linguistic task caused an increase, rather than decrease, in activity for both the hard (LH: *p*=0, *d*=0.74; RH: *p*=0, *d*=0.96) and easy (LH: *p*=0, *d*=0.79; RH: *p*=0, *d*=0.81) conditions. Furthermore, the RH fROI responded more strongly to the hard version than the easy version (*p*=0.001, *d*=0.45) and the LH fROI showed a similar trend (*p*=0.06, *d*=0.28). This effect was opposite in direction to the effect observed in language-responsive fROIs, indicating that the latter were functionally dissociable from other functional sub-regions in the hippocampus (LH: *p*<10^−12^, *d* = 1.04; RH: *p* <10^−7^ *d*=0.72).

### Experiment 2: Language-responsive fROIs are language-selective

In Experiment 2, we tested the recruitment of language-responsive fROIs during five non-linguistic tasks that placed high demands on executive processes like working memory and inhibitory control (*n*=11-15 participants in each task). To test whether these fROIs modulated their response with increasing cognitive demands, each task had an easy version and a hard version. The tasks (Fig. 1C) included: mental addition (“math”), a verbal working memory task (vWM), numeric and verbal multi-source interference tasks (nMSIT and vMSIT, respectively) and the Stroop task (56). Importantly, although some of these tasks use verbal materials, none of them require the binding of linguistic meanings to create more complex meaning representations, in contrast to the sentence comprehension condition; presumably, this is why these tasks do not activate the neocortical language regions (55; see also Discussion). If hippocampal language-responsive fROIs are selectively engaged in relational binding of linguistic representations (and not, e.g., numeric or spatial representations), then their engagement in these tasks should be weaker than their engagement during sentence comprehension̤

Consistent with this reasoning, our results (Fig. 3B) indicated that the recruitment of language-responsive fROIs, especially in the LH, was highly selective for linguistic processing. Descriptively, all non-linguistic tasks (except for the Stroop task in the RH fROI) led to deactivations that were not significantly below the fixation baseline (for all tasks, LH: *p*>0.09, 0<*d*<0.99; RH: *p*>0.12, 0.02<*d*<0.85). Therefore, these fROIs were not engaged in these non-linguistic tasks. Moreover, most deactivations during the hard versions of these tasks differed from the activations observed during the sentence reading condition of the language localizer, or showed a strong effect trending in that direction (LH: vWM, *p*=0.02, *d*=1.36; nMSIT, *p*=0.04, *d*=1.13; vMSIT, *p*=0.04, *d*=1.29; math: *p*=0.09, *d*=0.86; RH: vWM: *p*=0.02, *d*=2.01; nMSIT: *p*=0.01, *d*=1.36; vMSIT: *p*=0.07, *d*=1.08). The only tasks whose hard version did not significantly differ from sentence reading in their elicited responses were the Stroop task bilaterally (LH:*p*=0.14, *d*=0.73; RH: **p*=1*, *d*=0.03) and the math task in the RH (*p*=0.44, *d*=0.45). We note, however, that it is difficult to interpret such direct comparisons between response magnitudes due to many differences in design and procedure across tasks.

Bearing such differences in mind, a particularly interesting result is the deactivation observed during the hard version of the verbal working memory task, which mirrors the patterns previously observed for the neocortical language regions (55). This is an intriguing finding given that both tasks use verbal materials (lists of digit names or sentences, respectively) and engage relational processes (memorizing digit order or constructing a sentence-level meaning representation, respectively). Nonetheless, only the sentence comprehension task requires the encoding of relations of a linguistic (and hierarchical) nature among the words; the working memory task instead requires a (linear) combination of digits into a string, which could be performed over non-linguistic number (or other) representations. These results therefore suggest that the language-responsive fROIs may be specifically engaged in constructing complex linguistic meanings.

### Experiment 3: Activity in language-responsive fROIs is specifically correlated with activity in the neocortical language network

In Experiment 3, we evaluated the synchronization between hippocampal language-responsive fROIs and regions of the neocortical language network (localized with the reading task described above; 49). In other words, we tested whether the BOLD signal in hippocampal fROIs fluctuated over time in a similar pattern to the BOLD signal in neocortical fROIs. Specifically, we computed the correlations between the respective BOLD signal time-courses of these regions during each of two naturalistic conditions: “rest” (*n*=85), and auditory story comprehension (*n*=17) of narratives that are likely to place increased demands on hippocampal processing (e.g., through references to discourse entities introduced earlier in the story, 34, 35, 36) (see Supplemental Materials). The neocortical language network has been previously shown to exhibit strong synchronization among its regions during these conditions (59); if the language-responsive hippocampal fROIs are functionally integrated with this network, then they would similarly be functionally correlated with it.

Importantly, if the language-responsive hippocampal fROIs are functionally distinct from the rest of the hippocampus, then we should not observe these results for control fROIs. Moreover, if the engagement of these fROIs in linguistic processing is functionally specific, then their synchronization with the neocortical language network should not generalize to other neocortical networks. To assess this specificity, we also computed functional correlations between language-responsive hippocampal fROIs and the fronto-parietal “multiple demand (MD)” network (60, 61), a domain-general set of regions engaged in a wide variety of tasks, both linguistic and non-linguistic (localized with the spatial working memory task described above) (56, 62-66). Critically, the MD network does not show any synchronization with the neocortical language network during naturalistic cognition (59); if the language-responsive hippocampal fROIs are selectively integrated with the neocortical language network, then they should be dissociated from the MD network.

Overall, the hippocampal fROIs showed weak correlations with the neocortex, as is characteristic of subcortical structures (e.g., 67). Nonetheless, the pattern of correlations indicated a clear functional architecture consistent with our hypotheses (Fig. 3C), with strong effects especially in the story comprehension condition. In the LH, the language-responsive fROI was functionally correlated with the neocortical language network during both rest (mean Fisher-transformed correlation across neocortical regions and participants: *r*=0.08; *p*=0.004, *d*=0.57) and story comprehension (“stories”; *r*=0.15, *p*=0.004, *d*=1.24). These correlations showed two signatures of functional specificity. First, the neocortical language network was not functionally correlated with the control fROI (rest: *r*=0.02, *p*=1, *d*=0.11; stories: *r*=0.04, *p*=0.72, *d*=0.33), which thus differed from the language-responsive fROI (rest: *p*=0.01, *d*=0.36; stories: *p*=0.03, *d*=0.78). This result indicated that the neocortical language network was not uniformly correlated with the entire hippocampus. And second, the language-responsive fROI was uncorrelated (or even anti-correlated) with the neocortical MD network in the LH (rest: *r*=-0.06, *p*=0.01, *d*=0.4; stories: *r*=-0.04, *p*=0.52, *d*=0.52), a finding that distinguished this network from the neocortical language network (rest: *p*<10^−7^, *d*=0.72; stories: *p*<0.001, *d*=0.82). This result indicated that the language-responsive fROI was not uniformly correlated with the entire LH neocortex. Moreover, the language and MD networks significantly differed in that only the former was more correlated with the language-responsive fROI than with the control fROI (rest: *p*<10^−5^, *d*=0.59; stories: *p*<0.002, *d*=1.2).

Similar results obtained when testing correlations between (a) the LH hippocampus and the RH neocortex (rest: *p*<10-7, *d*=0.77; stories: *p*<0.001, *d*=1.33); and (b) the RH hippocampus and the RH neocortex (rest: *p*<0.002, *d*=0.51; stories: *p*=0.03, *d*=0.88). The correlation patterns between the RH hippocampus and the LH neocortex showed similar, but weaker, trends.

## Discussion

“The limbic cortex has no capacity for language”. This statement, from a TED talk viewed by nearly 30 million people (https://www.ted.com/talks/simon_sinek_how_great_leaders_inspire_action), reflects a popular view: language - a human artifact that is evolutionarily recent (43) - is processed exclusively by the neocortex and not by phylogenetically older structures. Contrary to this view, we provide evidence that the hippocampus is robustly and reliably recruited during language processing in healthy adults. This finding complements prior observations of hippocampal engagement in language tasks obtained with direct recordings of neural activity in patients with epilepsy (68–71). It is also consistent with neuropsychological findings from patients with hippocampal damage who show language processing impairments when relationships among linguistic elements need to be maintained and integrated into the unfolding representation of the discourse (6, 34-39).

Critically, to go beyond these prior findings, we used functional localization of language-responsive regions within the hippocampi of each participant in a large cohort and showed that such regions, especially in the left-hemisphere, exhibit a language-selective functional profile: they engage robustly and reliably during sentence comprehension, but are not engaged by six non-linguistic tasks even though these tasks require relational binding and/or include verbal materials. This region is also functionally synchronized in its activity fluctuations with the neocortical language network, but not with the domain-general multiple demand (MD) neocortical network. Furthermore, these functional signatures were spatially restricted to a portion of the hippocampus rather than being ubiquitously present throughout the hippocampus: we did not observe them in regions that were functionally localized with a non-linguistic, working memory task that required relational binding of spatial locations. Overall, our findings do not only indicate that the hippocampus “has a capacity for language” but also that this capacity, at least in part, relies on domain-specific rather than domain-general mechanisms (e.g., an “episodic buffer”; 72, 73).

This domain-specific contribution - observed here in adult brains - in no way entails that such specificity is present at birth (74, 75). Perhaps most plausibly, language-specific hippocampal mechanisms develop through our early, extensive experience with language. The localized implementation of these mechanisms in a focal sub-region also need not entail that they are innate, given that localized representations have well established metabolic, volumetric and computational advantages (76, 77). To go beyond such speculations in future work, it will be critical to track the developmental trajectory of this region during ontogeny, and relate it to the development of the neocortical language network with which it is selectively coupled.

What relational processes does the language-selective hippocampal region mediate? Preliminary evidence comes from an experiment reported in the Supplementary Materials, in which we found that this region was sensitive to “semantic composition” but not to “syntactic composition”. Namely, it responded more strongly to reading sentences, which allow for relational binding of word meanings, than to reading “Jabberwocky” sentences, which only allow for relational binding of abstract grammatical structures. In fact, the low response to reading “Jabberwocky” sentences was indistinguishable from the response to reading lists of either words or nonwords. This finding is inconsistent with earlier claims of syntactic integration in the hippocampus (71) (also see 41, 78), unless syntactic processing is construed as highly lexicalized; but it fits with a recent direct recording study of hippocampal activity demonstrating sensitivity to the semantic relatedness of words within a sentence (68), and with the deficits in word association tests observed in patients with hippocampal damage (79).

Still, the precise nature of hippocampal contributions to online language processing remains to be elucidated. At least three key questions are currently outstanding: first, is the domain-specific hippocampal region engaged in semantic processing of language per se or does it contribute to the integration of conceptual information more generally? For instance, would it be engaged in representing events that are depicted in non-linguistic, visual stimuli? Representing events and episodes has been linked to the hippocampus as a member of the Default Mode Network (DMN) (80, 81), a set of regions engaged in mind-wandering, self-referential thought, retrospection, prospection, and narrative processing (82–87); and the hippocampus in particular might also be the origin of spontaneously generated thoughts (88–90). Language, in turn, might be tightly linked to such “mental travel” into distant places, times, or minds and allows us to communicate such internal narratives to others; according to some, language has actually evolved for these purposes (91, 92).

Consistent with this link, the domain-specific hippocampal region exhibits some functional characteristics of the DMN, such as deactivation to non-linguistic tasks that require attention to external stimuli rather than to internal thoughts (93–96). Nevertheless, it shows some critically deviant characteristics such as activity increases during language comprehension (also see Supplementary Figure S1), thus combining functional signatures of both the DMN and language networks. It is therefore conceivable that the domain-specific hippocampal region is specialized for linguistic processing and not narrative thought in general, especially given that language can be functionally dissociated from such thought (e.g., representations of “who knows what” rely on non-linguistic mechanisms, and they remain largely intact despite hippocampal damage; 48, 97).

Second, we note that reliable and robust engagement during language processing was observed not only in the domain-specific hippocampal region but also in the control region that was localized for comparison purposes. However, these different regions exhibited distinct functional profiles; for instance, only the domain-specific region distinguished between sentences and nonword lists. We therefore argue that different hippocampal regions that are engaged in language processing implement distinct computations; but what is the division of labor between them, and is it organized along the same principles that govern the division of linguistic labor across neocortical networks? (98)

Third, what are the differences between hippocampal and neocortical contributions to language use? These contributions are clearly distinct, because patients with hippocampal damage perform well on many standard language tasks (16–18) whereas patients with damage to the cortical language regions exhibit aphasia. Our findings in the Supplementary Materials point to some initial differences between the domain-specific hippocampal region and the neocortical language network: whereas the former is mostly recruited during semantic composition, the latter is engaged in processing both lexical information and compositional syntax and semantics (49, 99, 100).

Given that hippocampal contributions to online language processing are less critical than those of the neocortical language regions, we do not consider it to be a “core” constituent of the language network. Indeed, not every brain region engaged during language processing is part of the core language network; for instance, the domain-general MD network is engaged by effortful comprehension (101) but is functionally dissociated from the language network (59). Critically, however, the hippocampal region we identified is functionally synchronized with the neocortical language network and appears to be similarly domain-specific. In light of this functional profile, we suggest to view this region as a peripheral node, or “satellite” member, of the language network. Expanding the language network in this way is a crucial step towards a more comprehensive understanding of the varied set of mechanisms that support online language processing.

## Methods

### Participants

One hundred and fifty participants (106 females) between the ages of 18 and 50, recruited from the MIT student body and the surrounding community, were paid $60 for their participation. All participants were right-handed native speakers of English, and naïve to the purposes of the experiments. They gave informed consent in accordance with the requirements of MIT’s Committee on the Use of Humans as Experimental Subjects (COUHES).

All participants completed the functional localizers used to define hippocampal fROIs. However, different, partially overlapping, subsets of our sample participated in different experiments: in Experiment 1, to test the size, robustness, and reliability of the localizer effects in the language-responsive and control fROIs, we used data from 90 participants (62 females) and 71 participants (50 females), respectively. In Experiment 2, to test the recruitment of the language-responsive fROIs during non-linguistic tasks, we used data from 11-15 participants for each task, with 36 participants (25 females) overall; and in Experiment 3, to test the functional synchronization between hippocampal and neocortical fROIs, we used data from 85 participants (57 females) in the resting-state condition and 17 participants (11 females) in the story comprehension condition.

### Design, materials and procedure

#### Language localizer

Language-responsive hippocampal fROIs were identified using a passive language comprehension task contrasting reading of sentences and lists of unconnected, pronounceable nonwords (49), a localizer task which has been previously shown to generalize across materials and modality of presentation (visual or auditory) (49, also see 102, 103). Sentence and nonword conditions were ordered in a standard blocked design with a counterbalanced order across runs (for timing parameters, see Supplementary Table S1). Stimuli were presented one word / nonword at a time. For about 75% of the participants, each trial ended with a memory probe and they had to indicate, via a button press, whether or not that probe had appeared in the preceding sentence / nonword sequence. Other participants simply read the materials passively and pressed a button at the end of each trial (a task that was added to maintain attention). Importantly, the sentences > nonwords effect of interest has been shown to generalize across such changes in task: regardless of whether the more engaging/demanding stimuli are sentences (in passive reading) or nonwords (in a memory task, because they are harder to memorize), the neocortical language network responds more strongly to sentences (49). Here, the same was true for the language-responsive hippocampal fROIs (see Supplementary Figure S1). Therefore, we report data collapsed across the two versions of the localizer task, which allows us to use a larger number of participants for functionally characterizing these fROIs. In addition, the size of the sentences > nonwords effect did not show sex differences in Experiment 1 (LH: *d*=0.3, *p*=0.15; RH: d~0, *p*~1), in line with the lack of such differences for neocortical language regions (104), so such differences were not explored further.

#### Non-linguistic localizer

For most participants, control fROIs in the hippocampus were localized with a spatial working memory task (55). On each trial, participants saw a 3*4 grid and kept track of four (easy condition) or eight (hard condition) locations that were sequentially flashed one or two at a time, respectively. Then, participants indicated their memory for these locations in a 2-alternative forced-choice (2AFC) paradigm, via a button press. Feedback was immediately provided upon choice (or lack thereof). Hard and easy conditions were presented in a standard blocked design with a counterbalanced order across runs (for timing parameters, see Supplementary Table S2). To localize hippocampal fROIs engaged in this task, we chose the hard > fixation contrast instead of the hard > easy contrast, as the latter did not give reliable results (both conditions require the relational binding of four sequentially presented visual displays and differ only in the number of locations per display; therefore, the demands on relational binding are perhaps too similar across conditions and their contrast might not be robust enough to detect hippocampal activity).

For participants in Experiment 2, who did not perform this task, control fROIs were instead localized with the contrast nonwords > fixation from the language localizer task (see 56; note that all of these participants performed the memory probe version of the task).

#### Experiment 2

Participants performed five non-linguistic tasks that engage working memory or cognitive control processes: mental arithmetic, a verbal working memory task, numeric and verbal multi-source interference tasks (105), and the Stroop task (for a brief description, see Supplementary Materials; for full details, see: 55).

#### Experiment 3

In the “rest” condition, participants were instructed to keep their eyes closed for 2-10min and let their mind wander. In the story comprehension condition, participants listened to 1-6 stories (see Supplementary Materials), each lasting 4.5-6min. These stories, based on publicly available texts, were edited to include linguistic phenomena that are infrequent in natural language; some of these phenomena (e.g., non-local syntactic dependencies) place high demands on flexible relational binding.

### Data acquisition and preprocessing

Structural and functional data were collected on a whole-body 3 Tesla Siemens Trio scanner with a 12-channel (n=32) or 32-channel (n=118) head coil at the Athinoula A. Martinos Imaging Center at the McGovern Institute for Brain Research at MIT. Acquisition parameters for these data varied slightly across participants scanned for different experiments (see Supplementary Materials). Preprocessing of functional data was carried out with SPM5 and included motion correction, normalization into a common space (Montreal Neurological Institute (MNI) template), resampling into 2mm isotropic voxels, smoothing with a 4mm FWHM Gaussian filter and high-pass filtering at 200s.

For Experiment 3, additional temporal preprocessing of the data was carried out using the CONN toolbox (106) in SPM8 as described in (59). Briefly, BOLD signal fluctuations originating from the white matter (WM) and cerebrospinal fluid (CSF) were regressed out of each voxel along with offline-estimated motion parameters, and the residuals were bandpass filtered to preserve only slow fluctuations (0.008-0.09 Hz).

### fROI definition

Language-responsive and control fROIs in the hippocampus were defined individually in each participant. For each localizer, a general linear model (GLM) estimated the effect size of each condition in each experimental run. These effects were each modeled with a boxcar function (representing entire blocks) convolved with the canonical Hemodynamic Response Function (HRF). The model also included first-order temporal derivatives of these effects, as well as nuisance regressors representing entire experimental runs and offline-estimated motion parameters. The obtained beta weights were then used to compute the functional contrast of interest: sentences > nonwords for language-responsive fROIs, and for control fROIs, either hard > fixation (in the non-linguistic task) or nonwords > fixation (in the reading task). Next, the resulting contrast maps were intersected with (a) an anatomical mask of the hippocampi (created with the wfu_pickatlas tool; 107); and (b) a binary mask of the participant’s gray matter (GM) based on SPM’s probabilistic segmentation of their structural data, where a voxel *v* was considered in the GM if: *p*(*v* ϵ **GM** >^max^ (*p*(*v* ϵ WM), *p(v* ϵ CSF)). Finally, in each hemisphere, the voxels falling within the hippocampus were sorted according to their localizer contrast values and the top 10% were defined as a fROI. This top n% approach ensures that fROIs can be defined in every participant and that their sizes are the same across participants, thus enabling us to generalize the results to the entire population (58).

For Experiment 3, fROIs in the neocortical language network were defined using group-constrained, participant-specific localization (49). fROIs in the neocortical MD network were defined using anatomically-constrained, participant-specific localization (56) (for a brief description, see Supplementary Materials).

### Characterizing fROI topography

The locations of fROIs should be characterized along spatial dimensions that carry relevant information about the shape of the hippocampus. To recover such dimensions, we applied principal components analysis to the three-dimensional coordinates of all hippocampal voxels, separately in each hemisphere. The first recovered axis explained over 78% of the variance in the spatial location of voxels, and correlated strongly with the MNI posterior-anterior axis (LH: *r* = 0.98; RH: *r* = 0.97) and inferior-superior axis (LH: *r* = −0.96; RH: *r* = −0.96), but weakly with the right-left axis (LH: *r* = 0.11; RH: *r* = −0.25). The second recovered axis explained over 17% of the variance, and correlated strongly with the MNI right-left axis (LH: *r* = 0.98; RH: *r* = 0.96), but weakly with the posterior-anterior axis (LH: *r* = 0.14; RH: *r* = 0.17) and the inferior-superior axis (LH: *r* = −0.16; RH: *r* = 0.15). The third axis explained less than 4% of the variance and was not considered further. The first two axes were then used as a coordinate system for describing the center of mass of each fROI in each participant.

### Data analysis

#### Estimating responses to the localizer tasks

Responses to both the language task and the non-linguistic task were estimated in each fROI using across-runs cross-validation (58). We used a subset of the runs (either odd or even) of a localizer task to define a fROI and the remaining runs (even or odd, respectively) to estimate the size of the localizer contrast effect in that fROI; the effect sizes so obtained were then averaged across the two estimations. This method ensures that the data used for fROI definition are independent of those used for response estimation and enables using all available data for both purposes (108, 109).

#### Experiment 2

Data from each task were analyzed using the GLM approach used for analyzing localizer data.

#### Experiment 3

Functional correlations between the hippocampus and neocortical networks were computed separately for each experimental condition. For each participant and fROI, BOLD signal time-courses were first temporally z-scored in each voxel and then averaged across voxels. Next, we computed Pearson’s moment correlation coefficient between every possible pair of time-courses consisting of one hippocampal fROI and one neocortical fROI. Correlations were Fisher-transformed to improve normality (110) and, finally, averaged across neocortical fROIs belonging to the same functional network and hemisphere. Thus, for each condition, we obtained 16 correlations: between the language-responsive / control hippocampal fROI in the left / right hemisphere and the neocortical language / MD network in the left / right hemisphere.

#### Statistical tests

Due to small sample sizes in some tasks and the resulting uncertainty regarding the normality of the data, we tested all planned comparisons using non-parametric statistics (in cases where samples were large, similar results were obtained using parametric statistics). Specifically, (i) condition means were compared to zero based on an empirical t-distribution generated from 9,999 bootstrap samples (in Experiment 3, similar results were obtained when the empirical null distribution of functional correlations was based on phase-randomized BOLD signal time-courses, following 111); (ii) all pairwise comparisons were tested using exact permutation tests for either two dependent or two independent samples (112); and (iii) 2 **x** 2 planned contrasts testing for region **x** condition interactions were converted to pairwise comparisons via their contrast weights (i.e., −1 and 1) and tested with the same exact permutation tests (113). For these comparisons of functional contrasts across regions, we accounted for regional differences in general responsiveness. Similarly, when comparing different functional contrasts within the same region, we accounted for task differences in general effectiveness (see Supplementary Materials). Within each experiment, all tests are presented following False Discovery Rate (FDR) correction for multiple comparisons (57).

## Supplementary Materials Methods

### Design for Experiment 2

In the mental addition task participants saw an integer and sequentially added three addends to it from either among the integers 2-4 (easy version) or 6-8 (hard version), and then chose the correct sum in a 2AFC paradigm. In the verbal working memory task, participants kept track of a sequence of four (easy) or eight (hard) single-digit number-words, and then chose the correct digit sequence in a 2AFC paradigm. In the numerical multi-source interference task, participants saw triplets of digits that included two occurrences of a certain digit and a single occurrence of a different digit, and pressed whichever button (from amongst a set) was labeled with the identity of the non-repeated digit; the position of this digit in the triplet either corresponded to the spatial position of its button on the button box (easy) or not (hard). In addition, the repeated digit either did not have a corresponding button (easy) or was a distractor from the button set (hard). In the verbal multi-source interference task, the digits were replaced with spatial words (“left”, “middle”, “right” or “none”). In the Stroop task, participants named the font-color of presented words, with the words themselves being either non-color adjectives (easy) or color adjectives different from the font color (hard).

### Data acquisition

Tl-weighted images were collected in 176 sagittal slices (1mm isotropic voxels; TR = 2,530ms; TE = 3.48ms). Functional, blood oxygenation level-dependent (BOLD) data were acquired using an EPI sequence with a 90^o^ flip angle and using GRAPPA with an acceleration factor of 2; the following parameters were used: thirty-one 4mm thick near-axial slices acquired in an interleaved order (with 10% distance factor), with an in-plane resolution of 2.1mm **x** 2.1mm, FoV in the phase encoding (A >> P) direction 200mm and matrix size 96mm **x** 96mm, TR = 2000ms and TE = 30ms. The first 10s of each run were excluded to allow for steady state magnetization.

### Defining fROIs in neocortical networks

fROIs in the neorocrtical language network were identified using group-constrained, participant-specific localization (49). For each participant, the sentences > nonwords contrast map was intersected with (a) a binary mask of the participant’s gray matter (see *Methods* section in the main text); and (b) binary masks based on a group-level representation of the sentences > nonwords contrast data obtained from a sample of 220 participants (partially overlapping with the sample reported here). The latter masks constrain the participant-specific fROIs to fall within neocortical areas in which activations for the localizer contrast are likely across the population. We used six such masks in the left hemisphere (LH), including regions in the posterior temporal lobe, anterior temporal lobe, angular gyrus, middle frontal gyrus (MFG), inferior frontal gyrus (IFG) and its orbital part. These masks were mirror-projected onto the right hemisphere (RH) to create six homologous masks (the masks cover significant parts of the cortex, so their mirrored version is likely to encompass the right-hemispheric homologue of the left-hemispheric language network, despite possible hemispheric asymmetries in their precise locations). These RH homologues were included because they are activated during many language-processing tasks (54, 114-117) and they show strong functional correlations with the LH language network during naturalistic cognition (59). In each of the resulting 12 masks, a fROI was defined using the top 10% method described in the *Methods* section of the main text.

fROIs in the neorocrtical MD network were identified similarly, but instead of using masks based on group-level data, we used anatomical masks (107) following (56). Nine masks were used in each hemisphere, including regions in the opercular IFG, MFG and its orbital part, insular cortex, precentral gyrus, Supplementary and preSupplementary motor area, inferior and superior parietal lobe, and anterior cingulate cortex.

### Comparing functional contrasts across regions and tasks

Each experimental task was designed to test one or more critical functional contrasts (e.g., sentences > nonwords in the language comprehension task). Such contrasts cannot be directly compared to one another across different regions or tasks, because any observed differences would be confounded with differences in general responsiveness or signal-to-noise ratio (118), e.g., due to vascularization (119–122). For instance, let *A, B* be regions that respond to sentences twice as strongly as they respond to nonwords. Here, there is a main effect of condition but, functionally, there is no region **x** condition interaction. However, assume further that region *A* responds more strongly than region *B* to language processing in general, such that there is a main effect of region. This main effect will make the sentences > nonwords contrast numerically bigger for region *A* than for region B, possibly driving the interaction towards significance (see 118). A similar scenario holds for the case where *A, B* are tasks testing different functional contrasts within a single region, with a main effect of task *A* engaging this region more strongly than task *B*. Therefore, prior to such tests in each experiment we pooled all functional contrasts, across all regions, into a single vector and regressed them against their respective markers of general responsiveness. Namely, the sentences > nonwords contrast was regressed against the response to sentences; and any contrast for a non-linguistic task was regressed against the response to the easy condition of that task. This regression was performed via a linear, mixed-effects model with a fixed effect for the general responsiveness markers and a random slope for participant, with no intercepts.

### Experiment S1: Language-responsive fROIs in the LH are sensitive to semantic composition

To further functionally characterize the contributions of language-responsive hippocampal fROIs to online language use, we asked which aspects of language processing recruited them. Therefore, we tested their sensitivity to lexical (i.e., word-level) information and compositional (i.e., sentence-level) information. For this purpose, we measured fROI responses in 31 participants (20 females) to reading (i) lists of unrelated words, involving lexical but not compositional processing; (ii) “Jabberwocky” sentences in which content words are replaced by nonwords, involving compositional processing of abstract structure but no lexical processing; as well as (iii) sentences, involving both lexical and compositional processing, and (iv) lists of nonwords, involving neither lexical nor compositional processing (Figure S2A). For a full description of the materials, as well as information regarding timing parameters, run structure and acquisition protocol, see Experiments 1&2 in (49).

Given the role of the hippocampus in relational binding, we expected the language-responsive fROIs to increase their activity in the presence of compositional information and thus contrasted two hypotheses: (i) language-responsive fROIs engage in composition regardless of whether semantically-specified lexical information is present or not, i.e., they compute relations between abstract grammatical categories and structures. In this case, we expect only a main effect of compositionality, such that both sentences and Jabberwocky engage these fROIs more strongly than word and nonword lists (sentences ~ Jabberwocky > word lists ~ nonword lists); (ii) language-responsive fROIs engage in composition only for semantically specified lexical units, i.e., they compute relations between real words. In this case, we expect an interaction between compositionality and lexical information, such that sentences engage these fROIs more strongly than Jabberwocky, but word and nonword lists both engage them weakly (sentence > Jabberwocky ~ word lists ~ nonword lists).

In the LH language-responsive fROI we found support for the second hypothesis, with an interaction between lexical and compositional processing (*p*=0.04, *d*=0.56). Specifically, this region responded more strongly to sentences than to Jabberwocky (*p*=0.002, *d*=0.81) but did not respond differentially to word-lists and nonwords (*p*=1, *d*=0.08). In other words, this fROI engaged in reading more strongly when word meanings were present compared to when they were absent, but only if such lexical information was embedded in a sentential context allowing for semantic composition. In the RH, too, responses to word-lists and nonwords did not differ (*p*=1, *d*=0.08), and there was a trend towards a moderate Sentences > Jabberwocky effect (*p*=0.06, *d*=0.51). Here, the interaction between lexical and compositional processing was not significant (*p*=0.41, *d*=0.31); nonetheless, we could not conclude that the two hemispheres differed in the size of this interaction (*p*=0.29, *d*=0.35) (Figure S2B).

This pattern of results was unique to the language-responsive fROIs: control fROIs strongly engaged in all kinds of reading (for all conditions, LH: *p*=0, *d*>0.98; RH: *p*=0, *d*>0.82) and indistinguishably so (for all pairwise comparisons, LH: *p*=1, *d*<0.15; RH: *p*=1, *d*<0.12). Therefore, compared to language-responsive fROIs, the ipsilateral control fROIs showed weaker effects for the Sentences > Jabberwocky contrast (LH: *p*<10^−5^, *d*=1.17; RH: *p*<0.001, *d*=0.81) as well as the Sentences > Nonwords contrast (LH: *p*=0.007, *d*=0.65; RH: *p*=0.007, *d*=0.67). Furthermore, in the LH, language-responsive and control fROIs differed in the sizes of the Sentences > Word-lists effect (*p*=0.009, *d*=0.67) and Word-lists > Jabberwocky effect (*p*=0.007, *d*=0.65). Critically, the interaction between lexical and compositional processing was weaker in the control fROIs compared to the ipsilateral language-responsive fROIs (LH: *p*=0.004, *d*=0.74; RH: *p*=0.048, *d*=0.53) (Figure S2B).

### Sample story used in Experiment 3

At ten years old, I could not figure out what it was that this Elvis Presley guy had that the rest of us boys did not have. He seemed to be no different from the rest of us. He was simply a man who had a head, two arms and two legs. It must have been something pretty superlative that he had hidden away, because he had every young girl at the orphanage wrapped around his little finger.

At about nine o’clock on Saturday morning, I figured a good solution was to ask Eugene Correthers, who was one of the older and smarter boys, what it was that made this Elvis guy so special. He told me that it was not anything about Elvis’s personality, but his wavy hair, and the way he moved his body. About a half an hour later, the boys in the orphanage called down to the main dining room by the matron were told that they were all going to downtown Jacksonville, Florida to get a new pair of Buster Brown shoes and a haircut. That is when I got this big idea, which hit me like a ton of bricks. If the Elvis haircut was the big secret, then Elvis’s haircut I was going to get.

I was going to have my day in the sun, and all the way to town that was all I talked about. The fact that I was getting an Elvis haircut, not just the simple fact that we were getting out of the orphanage, made me particularly loquacious. I told everybody, including the orphanage matron I normally feared, that I was going to look just like Elvis Presley and that I would learn to move around just like he did and that I would be rich and famous one day, just like him. The matron understood my idea was something that I was really excited about and said nothing.

When I got my new Buster Brown shoes, I was smiling from ear to ear. Those shoes, they shined really brightly, and I liked looking at the bones in my feet, which I had never seen before, through a special x-ray machine they had in the shoe store that made the bones in your feet look green. I was now almost ready to go back to the orphanage and practice being like the man who all the girls loved, since I had my new Buster Brown shoes. It was the new haircut, though, that I needed to complete my new look.

We finally arrived at the unassuming, unembellished barbershop, where they cut our hair for free because we were orphans. Even though we were supposed to slowly wait to be called, I ran straight up to one of the barber chairs and climbed up onto the board the barber placed across the arms to make me sit up higher. I looked at the man and said, with a beaming smile on my face, “I want an Elvis haircut. Can you make my hair like Elvis’s?” I asked. The barber, who was a genial young man, grinned back at me and said that he would try his best.

I was so happy when he started to cut my hair, but just as he started to cut, the matron, who had been watching me and had a look as cold as ice, motioned for him to come over to where she was standing. She whispered something into his ear that caused the barber to shake his head, like he was telling her, “No”. In response, the matron walked over to a little man sitting in an office chair that squeaked as it rolled around the floor and spoke to him. It was the little man who then walked over and said something to the man who was cutting my hair. The next thing I knew, the man who was cutting my hair told me that he was no longer allowed to give me an Elvis cut.

“Why not?” I cried desperately.

The kindly barber stopped by the matron did not answer, but from his expression, I could tell that he wished he could cut it as I had asked.

Within a few minutes, it wasn’t an Elvis haircut, but a short buzz cut that the barber had given me. When he finished shaving off all my hair and made me smell real good with his powder, the barber handed me a nickel and told me to go outside to the snack machine and buy myself a candy bar. I handed him the nickel back and told him that I was not hungry. “I’m so sorry, baby,” he said, as I climbed out of his barber chair. “I am not a baby,” I said, as I wiped the tears from my eyes.

I then sat down on the floor and brushed away the hair that had accumulated on my shiny new Buster Brown shoes. My head was no longer in the clouds, and I got up off the floor, brushed off my short pants, and walked sullenly towards the door.

The matron was smiling at me sort of funny like.

The barber upset by the matron said to her, “You are just a damn bitch, lady.”

She yelled back at him at the top of her lungs, before walking toward the office, as fast as she could.

To show his anger, the man hit the wall with his hand and then walked outside where he stood against the brick wall, smoking a cigarette. I understood right there my haircut was something that had been out of the power of the barber and then I slowly walked outside to join the man. He looked down, smiled at me, then he patted me on the top of my bald as a coot head. It was a fact of my life that I was not gonna have hair that was anything like Elvis’s anytime soon. I then looked up at the barber with my wet red eyes and asked, “Do you know if Elvis Presley has green bones?”

## ACKNOWLEDGEMENTS

Support for this research and manuscript provided by NICHD R00 HD057522 to EF and NIDCD R01 DC011755 to MCD and SBS. We would like to acknowledge the Athinoula A. Martinos Imaging Center at McGovern Institute for Brain Research at MIT, and the support team (Steve Shannon, Atsushi Takahashi, and Sheeba Arnold).

---

**Table S1.** Task design and timing parameters for different versions of the language localizer task

|  | Version |  |  |  |  |  |  |
| --- | --- | --- | --- | --- | --- | --- | --- |
|  | 1 | 2 | 3 | 4 | 5 | 6 | 7 |
| Number of participants<sup>a</sup> | 40 | 4 | 3 | 44 | 42 | 22 | 2 |
| Task: Passive Reading or Memory? | PR | PR | PR | M | M | M | M |
| Words / nonwords per trial | 12 | 12 | 12 | 12 | 8 | 8 | 8 |
| Trial duration (ms) | 6,000 | 3,600 | 4,800 | 6,000 | 4,800 | 4,800 | 4,800 |
| Fixation | 100 | 600 | 600 | --- | --- | --- | --- |
| Presentation of each word / nonword | 450 | 250 | 350 | 350 | 350 | 350 | 350 |
| Fixation | 500 | --- | --- | 300 | 300 | 300 | 300 |
| Memory probe | --- | --- | --- | 1,000 | 1,350 | 1,350 | 1,350 |
| Fixation | --- | --- | --- | 500 | 350 | 350 | 350 |
| Trials per block | 3 | 5 | 5 | 3 | 5 | 5 | 5 |
| Block duration (s) | 18 | 18 | 24 | 18 | 24 | 24 | 24 |
| Blocks per condition (per run) | 8 | 4 | 4 | 6 | 4 | 4 | 4 |
| Conditions<sup>b</sup> | S,N | S,W,J,N | S,W,J,N | S,W,N | S,W,N | S,W,J,N | S,W,J,N |
| Fixation block duration (s) | 14 | 18 | 24 | 18 | 16 | 16 | 24 |
| Number of fixation blocks | 5 | 5 | 5 | 4 | 3 | 5 | 5 |
| Total run time (s) | 358 | 378 | 504 | 396 | 336 | 464 | 504 |
| Number of runs | 2 | 8 | 7-8 | 2-3 | 3 | 4-8 | 7-8 |

<sup>a</sup> Participants who participated in more than one experiment were sometimes scanned on multiple days and performed different versions of the localizer task on different sessions.
<sup>b</sup> S = Sentences, N = Nonwords, W = Word lists, J = Jabberwocky sentences

**Table S2.** Task design and timing parameters for different versions of the non-linguistic task (spatial working memory) used to localizer control fROIs

|  | Version |  |  |
| --- | --- | --- | --- |
|  | 1 | 2 | 3 |
| Number of participants | 69 | 18 | 3 |
| Trial duration (ms) | 8,000 | 8,500 | 9,000 |
| Fixation | 500 | 500 | 500 |
| Sequential presentation of locations<sup>a</sup> | 4,000 | 4,000 | 4,000 |
| 2AFC | 3,250 | 3,750 | 4,250 |
| Feedback | 250 | 250 | 250 |
| Trials per block | 4 | 4 | 2 |
| Block duration (s) | 32 | 34 | 18 |
| Blocks per condition (per run) | 6 | 5 | 6 |
| Conditions | Easy, Hard | Easy, Hard | Easy, Hard |
| Fixation block duration (s) | 14 | 14\16<sup>b</sup> | 18 |
| Number of fixation blocks | 5 | 2\4 | 4 |
| Total run time (s) | 358 | 432 | 288 |
| Number of runs | 2-3 | 2-3 | 2 |

<sup>a</sup> In the easy condition, 4 single locations were sequentially presented. In the hard condition, 4 location pairs were sequentially presented.
<sup>b</sup> Pre-run and post-run fixation blocks were shorter than fixation blocks within the run.

**Figure S1.**
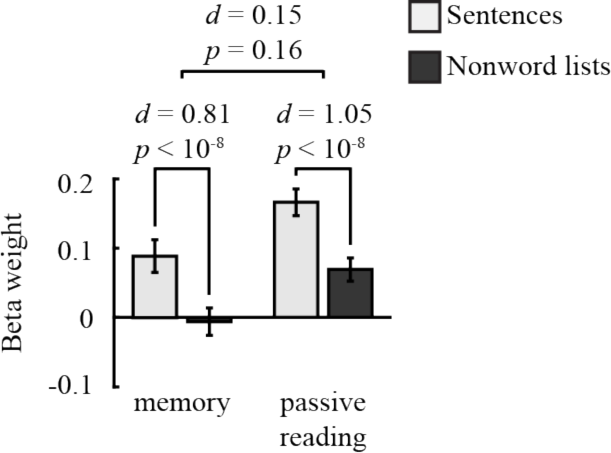
Effect sizes for two versions of the language localizer task in the left-hemisphere, language-responsive hippocampal fROI. Note that whereas the passive reading version appears to engage the fROI to a greater extent compared to the memory version for both sentences and non-words, the size of the sentences > nonwords effect is indistinguishable across the two versions.

**Figure S2.**
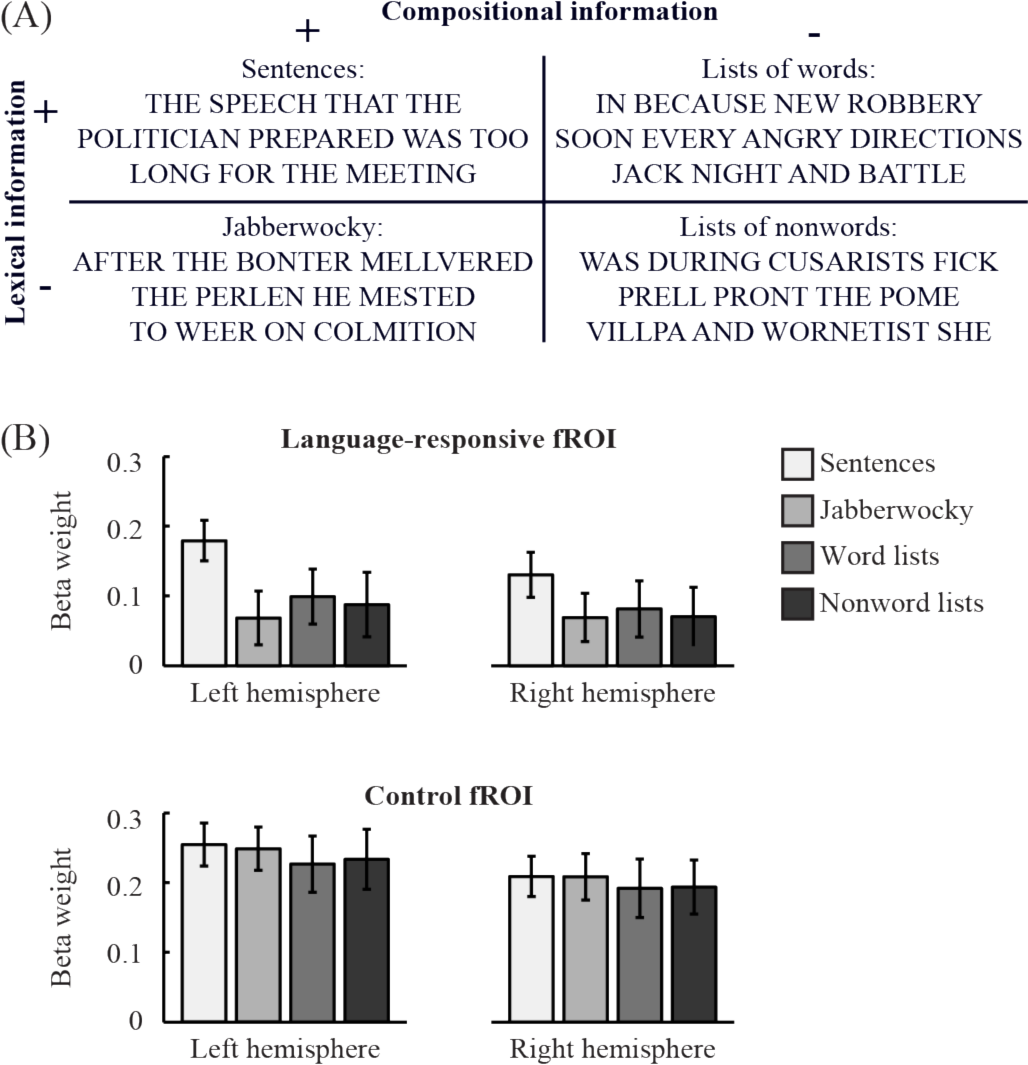
Sensitivity of hippocampal fROIs to lexical and compositional information. (A) Example for each stimulus category used in Experiment S1. (B) Effect sizes in each hippocampal fROI for each stimulus category.

